# A Refined Developmental Transcriptomic Landscape Reveals Four Coordinated Programs in *Chlamydia trachomatis*

**DOI:** 10.64898/2026.09.15.751914

**Authors:** Danny Wan, Wurihan Wurihan, Zhao Lai, Guangming Zhong, Huizhou Fan

## Abstract

The developmental cycle of *Chlamydia trachomatis* requires coordinated transitions between infectious elementary bodies (EBs) and replicative reticulate bodies (RBs), yet the temporal organization of the underlying transcriptional programs remains incompletely resolved. We performed high-coverage RNA sequencing from 0 to 32 h post-infection (hpi), with closely spaced early sampling, and used unsupervised clustering of temporal expression profiles to identify four coordinated programs: immediate-early host adaptation and developmental program, RB formation and maintenance, RB proliferation and secondary differentiation, and EB formation and invasion. The immediate-early program peaked at 1 hpi and contained a strong representation of inclusion-membrane and intracellular-trafficking genes, together with the developmental regulator gene euo. The RB formation and maintenance program peaked at approximately 3 hpi and sustained expression throughout much of the cycle, encompassing genes that support transcription, translation, and core metabolism. The RB proliferation and secondary differentiation program combined DNA replication and cell-division genes with alternative sigma factors, partner-switching components, and type III secretion apparatus genes, consistent with developmental preparation during proliferation. The EB formation and invasion program contained genes associated with structural maturation, host-cell interaction, and infectious competence. These findings distinguish transient host-adaptation expression from sustained early biosynthetic expression, separate RB establishment from population expansion, and differentiate regulatory preparation for the RB-to-EB transition from later EB maturation. Together, the four programs define partially overlapping transcriptional priorities that provide a framework for investigating how gene expression coordinates chlamydial growth, differentiation, and preparation for the next infection cycle.

## Introduction

*Chlamydia trachomatis* is an obligate intracellular bacterial pathogen responsible for a wide spectrum of human diseases, including trachoma and sexually transmitted infections (1). A hallmark of *Chlamydia* biology is a highly specialized biphasic developmental cycle involving two morphologically and functionally distinct forms: the infectious elementary body (EB) and the replicative reticulate body (RB) (2–4). EBs are metabolically limited but structurally adapted for extracellular survival and host cell invasion, whereas RBs are transcriptionally active and undergo proliferation within a membrane-bound inclusion, where they subsequently differentiate back into EBs prior to release from the host cell (2–4).

Following host-cell entry, EBs activate bacterial metabolism and undergo primary differentiation into RBs, reaching a midpoint at approximately 3 h post-infection (hpi) and completing by approximately 6-8 hpi (5). Proliferating RBs then replicate within the inclusion before undergoing secondary differentiation into EBs (2–4). Although the morphological progression of this cycle is well characterized, the transcriptional programs underlying these transitions remain incompletely resolved.

Early transcriptional profiling studies using DNA microarrays demonstrated that chlamydial gene expression is temporally regulated (5, 6). Belland et al. identified subsets of immediate-early and late genes associated with primary and secondary differentiation, respectively, and showed that transcripts of late genes are present in EBs prior to infection, suggesting transcript preloading (5). Nicholson et al. extended these observations by identifying multiple temporally coordinated gene clusters and proposing global transcriptional stages linked to developmental progression (6). However, in that study, the earliest sampling point was 6 hpi, a time at which EB-to-RB differentiation is largely underway or nearing completion, potentially limiting resolution of the earliest transcriptional events.

The advent of RNA sequencing (RNA-seq) has enabled more sensitive and comprehensive profiling of the chlamydial transcriptome (7). Recent high-MOI RNA-seq analyses revealed that transcriptional activation during the first hour post-infection is far more extensive than previously appreciated, with over 70% of the genome showing increased expression (8). These findings indicate that early differentiation is accompanied by widespread transcriptional activation rather than being driven by a limited set of immediate-early genes. However, whether these early-induced genes constitute a single transcriptional response or multiple temporally distinct programs has remained unclear.

To address this gap, we performed high-coverage RNA-seq across 0 to 32 hpi, using operationally optimized MOIs: higher MOIs at early time points to improve detection of bacterial transcripts and a lower MOI at later time points to limit premature host-cell lysis at high bacterial burdens (5). The resulting dataset enabled robust quantification of transcript abundance as transcripts per million (TPM) across the developmental cycle and defined a refined developmental transcriptomic landscape of *C. trachomatis*. Unsupervised clustering identified four temporally coordinated transcriptional programs: immediate-early host adaptation and developmental program, RB formation and maintenance, RB proliferation and secondary differentiation, and EB formation and invasion. These labels summarize the dominant functional themes associated with each coexpression trajectory; they do not imply that every gene within a cluster is dedicated exclusively to a single developmental process.

## Results

### High-coverage transcriptomic data reveal a coherent developmental trajectory across the *C. trachomatis* developmental cycle

To capture transcriptional dynamics across the *C. trachomatis* developmental cycle, samples were collected at time points spanning entry, primary differentiation, RB expansion, and late-stage development (0–32 hpi) (Fig. 1A). Early time points (0–3 hpi) were selected to capture events immediately following host cell entry and initiation of EB-to-RB differentiation, whereas mid-cycle (8–16 hpi) and late-stage (24–32 hpi) samples correspond to RB establishment, replication, and subsequent differentiation into EBs.

**Fig. 1.**
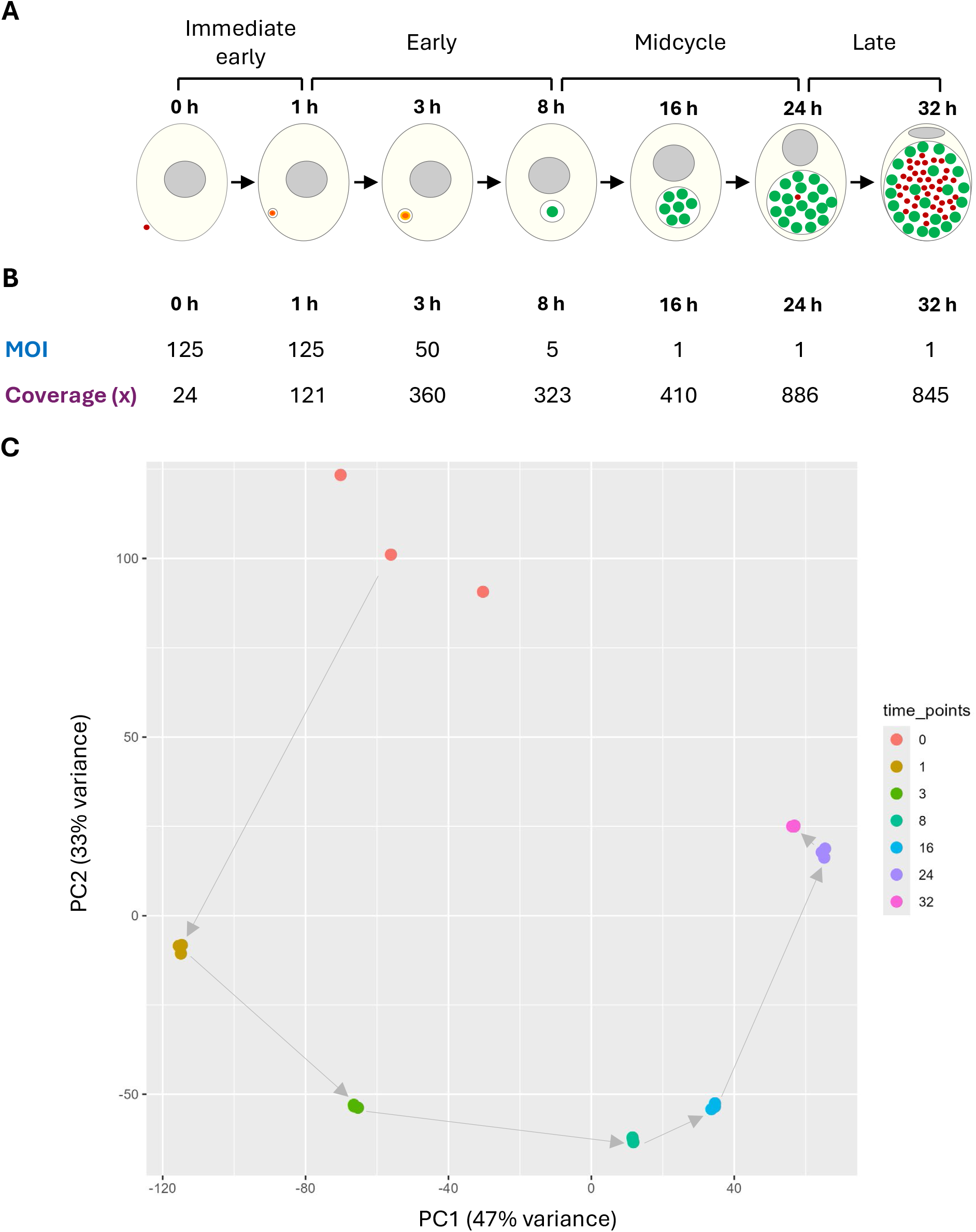
Experimental design and global transcriptomic organization of the *C. trachomatis* developmental cycle. (A) Schematic of the *C. trachomatis* developmental cycle and sampling strategy. Samples were collected at 0, 1, 3, 8, 16, 24, and 32 h post-infection (hpi), spanning entry, primary differentiation, RB proliferation, and late-stage development. (B) Multiplicity of infection (MOI) and transcriptome coverage across developmental stages. Higher MOIs were used at early time points to maximize recovery of chlamydial transcripts when bacterial RNA abundance is low, whereas lower MOIs were used at later stages to limit premature host-cell lysis at high bacterial burdens (5). At both 0 and 1 hpi, read counts from datasets generated at MOIs of 50 and 200 were pooled (8). The arithmetic mean of these MOIs is 125; this value does not represent a separately tested infection condition. Coverage values indicate average genome coverage at each time point. (C) Principal component analysis (PCA) of transcriptomes across the developmental cycle. Samples form a continuous developmental trajectory from early to late stages. Arrows indicate the direction of developmental progression.

To ensure robust detection of chlamydial transcripts across developmental stages, read counts from two previously reported RNA-seq datasets generated at MOIs of 50 and 200 were pooled for each of the 0- and 1-hpi time points (8). Progressively lower MOIs were used for subsequent time points (Fig. 1B). This strategy increased early transcript recovery while limiting premature host-cell lysis at high bacterial burdens during later stages. Consistent with this design, high genome coverage was achieved across all time points (Fig. 1B), enabling comprehensive measurement of chlamydial transcriptional dynamics across the developmental cycle.

To evaluate relationships among transcriptome-wide expression profiles across time points, principal component analysis (PCA) was performed on normalized gene expression values across all samples (Fig. 1C). The PCA revealed a coherent developmental trajectory, with samples forming a continuous progression from early to late stages. Early time points (0–3 hpi) were clearly separated from mid-cycle samples (8–16 hpi), which in turn were distinct from late-stage samples (24–32 hpi), indicating substantial global transcriptional remodeling over the course of infection.

Notably, the directionality of the trajectory mirrors the developmental progression depicted in Fig. 1A, with samples transitioning from entry and early differentiation to RB expansion and, ultimately, to late-stage development. This concordance between biological staging and transcriptomic organization indicates that the dataset captures coordinated, stage-associated transcriptional changes across the developmental cycle. Together, the high coverage and coherent global structure of the data provide a robust foundation for defining discrete transcriptional programs underlying *C. trachomatis* development.

### Identification of four transcriptional programs and their functional specialization across the developmental cycle

To define the temporal organization of gene expression across the developmental cycle, stage-specific TPM profiles were used to cluster gene-expression trajectories with an unsupervised approach. TPM values provided a common scale for comparing relative transcript abundance across time points despite differences in bacterial burden and for identifying coordinated expression trajectories (7).

The number of clusters was determined directly from the data using unsupervised clustering and the elbow method (9). The resulting plot showed a clear inflection at four clusters (Fig. 2A), supporting four as the best-supported representation of the principal transcriptional trajectories.

**Fig. 2.**
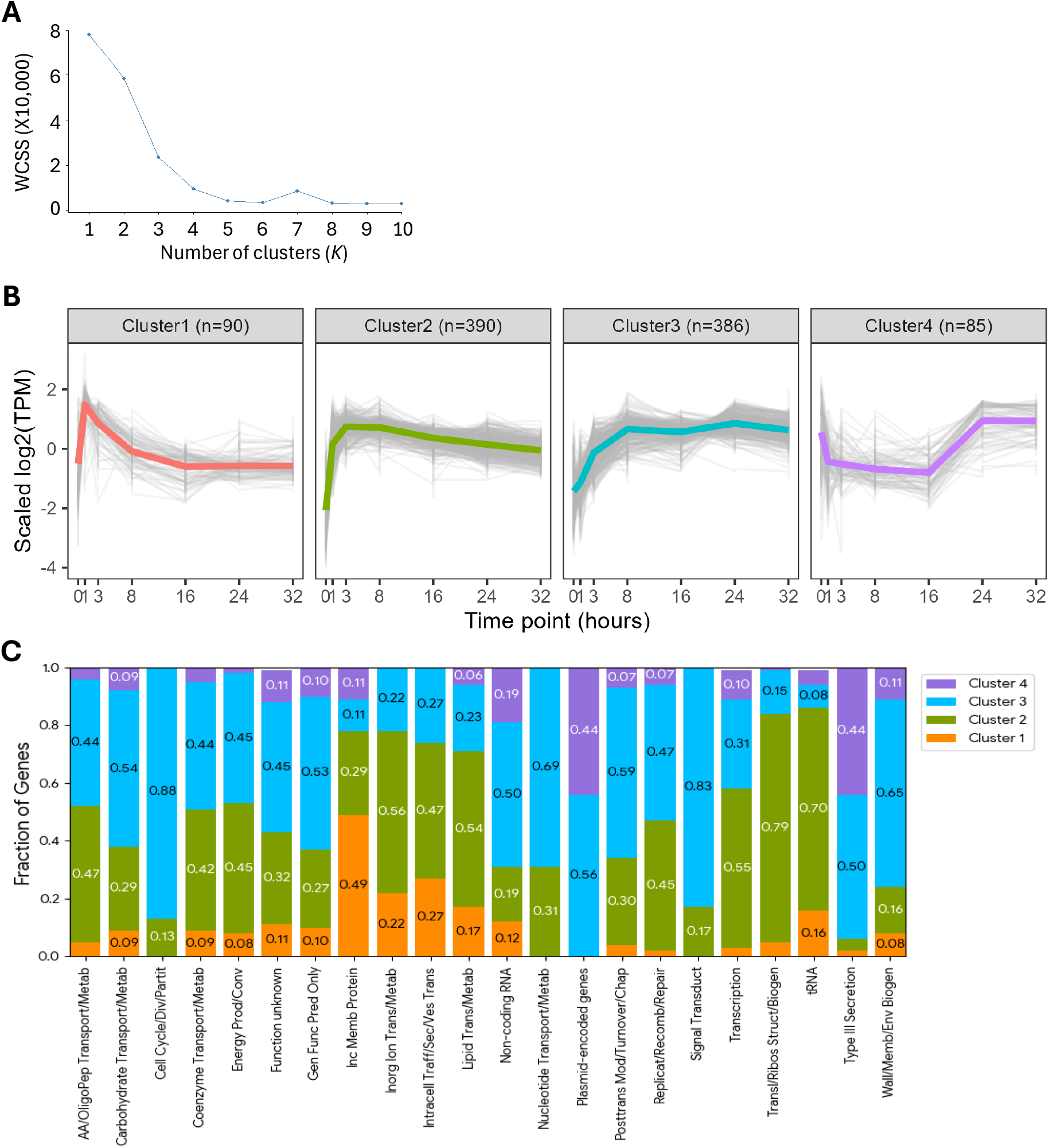
Identification of four coordinated transcriptional programs across the *C. trachomatis* developmental cycle. (A) Determination of the optimal number of clusters using the elbow method. The within-cluster sum of squares (WCSS) was calculated for increasing cluster numbers (*K*). The inflection point at K = 4 was selected for downstream analyses. (B) Temporal expression profiles of genes clustered into the four transcriptional programs. Gene expression values were normalized and clustered based on temporal trajectories across the developmental cycle. Shaded regions indicate the distribution of gene expression patterns within each cluster, and solid lines indicate cluster centroids. Cluster 1 represents an immediate-early host adaptation and developmental program, Cluster 2 an RB formation and maintenance program, Cluster 3 an RB proliferation and secondary differentiation program, and Cluster 4 an EB formation and invasion program. (C) Functional distribution of selected COG categories among the four transcriptional programs. Genes were classified into functional classes and the fraction of genes within each category represented in each cluster was determined. Colors correspond to the four transcriptional programs shown in panel B.

After exclusion of the six rRNA genes, 951 of the 957 annotated genes were included in clustering. The four clusters comprised 90, 390, 386, and 85 genes, representing 9.4%, 40.8%, 40.3%, and 8.9% of all annotated genes, respectively (Fig. 2B). Together, they define four temporally coordinated transcriptional programs underlying the developmental transcriptomic landscape of *C. trachomatis*. The complete gene membership and functional annotations for each cluster are provided in Tables S1-S4.

To characterize the biological content of each trajectory, genes within selected COG classes were further classified according to functional role, and their distribution across clusters was quantified (Fig. 2C). The clusters are defined by shared temporal expression patterns; accordingly, the interpretations below focus on the dominant functional themes and selected developmental regulators and pathway components represented in each trajectory.

#### Cluster 1: Immediate-early host adaptation and developmental program

Cluster 1 genes exhibited rapid induction following host-cell entry, peaked at 1 hpi, and declined progressively thereafter (Fig. 2B), defining a transient immediate-early program. Its functional composition indicates that host adaptation is the dominant biological theme: 17 of the 35 (49%) inclusion-membrane genes and 4 of the 15 (27%) genes involved in intracellular trafficking, secretion, or vesicle transport fell into Cluster 1 (Fig. 2C; Table S1). Inclusion membrane proteins form a critical host–pathogen interface and can modulate host pathways that support inclusion development and acquisition of host-derived nutrients (10–13). Their overrepresentation in Cluster 1 is consistent with rapid establishment of an intracellular niche that supports chlamydial development. Cluster 1 also contained *euo*, the only curated developmental regulator in this cluster. Although the developmental role of Cluster 1 is not fully apparent from the broad functional distribution in Fig. 2C, *euo* encodes a regulator that represses transcription of numerous late genes, indicating that early developmental regulation occurs alongside host adaptation (14–17).

#### Cluster 2: RB formation and maintenance program

Cluster 2 genes were rapidly induced between 0 and 1 hpi, reached peak expression at approximately 3 hpi, and then gradually declined at 8, 16, 24, and 32 hpi, yet remained substantially above their level at 0 hpi (Fig. 2B). This trajectory is consistent with a program initiated early and maintained throughout much of the developmental cycle. Functional analysis showed that Cluster 2 was dominated by genes involved in core biosynthetic and metabolic activities (Fig. 2C, Table S2). Notably, 55% of transcription-related genes, 79% of translation and ribosomal-structure genes, and 70% of tRNA-associated genes fell into this cluster. Substantial fractions of genes involved in inorganic-ion transport and metabolism (56%) and lipid transport and metabolism (54%) were also represented. The coordinated enrichment of transcriptional machinery, translational components, and tRNAs indicates sustained gene-expression capacity, while the representation of ion-transport and lipid-metabolism genes is consistent with continued metabolic and membrane homeostasis. The cluster also contained inclusion-membrane and secretory genes, consistent with continued development of the intracellular niche. Together, these functions support the designation RB formation and maintenance program.

The early peak of Cluster 2 expression at approximately 3 hpi may indicate that transcriptional, translational, and metabolic activities are particularly elevated during the period preceding completion of RB formation. Once RBs are established, the gradual decline in Cluster 2 expression may reflect reduced demand for these activities as the organisms transition into a sustained proliferative state.

#### Cluster 3: RB proliferation and secondary differentiation program

Cluster 3 genes increased sharply during the early phase of infection, reached high expression by approximately 8 hpi, and remained elevated through 32 hpi (Fig. 2B). They were strongly enriched for functions associated with chromosome replication, cell division, chromosome partitioning, nucleotide metabolism, and cell-envelope biogenesis (Fig. 2C, Table S3). These features identify RB proliferation as the dominant functional theme of the cluster.

Cluster 3 also contained 24 of the 48 T3SS genes distributed across the four clusters, including genes associated with the RB-expressed *ctl0238*, *copB2*, and *copD2* apparatus (Table S3). In addition, the cluster contained the alternative sigma factors *fliA* (σ28) and *rpoN* (σ54), as well as multiple signal-transduction genes, including components of partner-switching systems. The coexpression of these regulatory factors with replication, cell division, and T3SS structural genes provides a basis for linking RB proliferation to progression toward secondary differentiation. The replication and cell-division genes support the expansion of the RB population, whereas genes encoding alternative sigma factors (σ28 and σ54), components of partner-switching mechanisms, and T3SS structural apparatus are consistent with the acquisition of regulatory and functional capacities associated with secondary differentiation. Here, secondary differentiation refers primarily to the functional and regulatory reprogramming that prepares proliferating RBs for the subsequent RB-to-EB developmental transition. Together, these features support the designation RB proliferation and secondary differentiation program.

#### Cluster 4: EB formation and invasion program

Cluster 4 genes showed relatively high expression at 0 hpi, declined sharply after host-cell entry, remained at low levels during the early and mid-cycle stages, and were strongly induced again at 24–32 hpi (Fig. 2B). The 0-hpi signal is consistent with the presence of these transcripts in incoming EBs, whereas their late induction reflects renewed expression during EB maturation (Table S4). The cluster contained genes associated with EB structural maturation, including genes encoding the histone-like proteins HctA and HctB and the EB-associated outer-membrane proteins OmcA and OmcB (Table S4). It also contained genes associated with host-cell interaction, including the EB-associated T3SS genes *scc2*, *copB*, and *copD*, genes encoding numerous T3SS effectors, *tarp*, and four plasmid-encoded genes, including pgp3, which encodes a secreted virulence factor, and pgp4, which encodes a regulator of chromosomal gene expression (Fig. 2C, Table S4). These features distinguish Cluster 4 from Cluster 3, which contains many T3SS structural genes during RB proliferation and secondary differentiation.

The coordinated expression of EB structural proteins, T3SS components and effectors, and plasmid-encoded virulence-associated factors suggests that Cluster 4 supports the ordered late events of EB maturation, host-cell exit, and preparation for invasion in the next developmental cycle. Recent evidence that distinct T3SS translocons and effectors are expressed in different chlamydial cell forms supports the association of these late genes with EB entry competence (18). Together, these features support the designation EB formation and invasion program.

### Subcluster analysis reveals developmental shifts in metabolic strategies

Because broad COG categories often include genes with diverse physiological functions, selected metabolic gene groups were further subdivided according to their predicted metabolic roles (Fig. 3A–C). Among genes involved in energy production and conversion (Fig. 3A), all 3 of 3 Cluster

- genes fell into the energy-acquisition category. In contrast, energy generation dominated Clusters
- and 3, accounting for 14 of 16 genes (88%) and 15 of 17 genes (88%), respectively, whereas acquisition declined to 1 of 16 genes (6%) in Cluster 2 and was absent thereafter. The two Cluster 4 genes also fell into the energy-generation category. These data suggest a shift in energy-related transcriptional priorities during RB establishment, from energy acquisition toward endogenous energy generation.

**Fig. 3.**
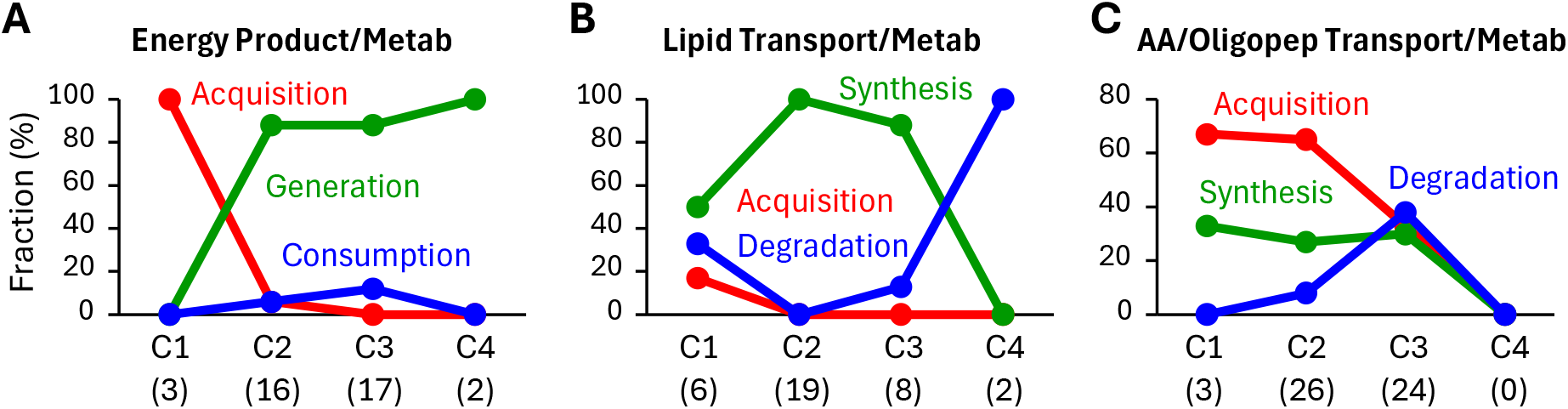
Developmental redistribution of metabolic functions across the four transcriptional programs. Selected metabolic genes were categorized according to predicted physiological roles, and their distribution across the four transcriptional programs was determined. Numbers in parentheses indicate the total number of genes represented in the corresponding cluster for each analysis. (A) Energy-production and conversion functions were most prominent in Clusters 2 and 3, whereas energy-acquisition functions were concentrated in Cluster 1. (B) Lipid-synthesis functions predominated in Clusters 2 and 3, whereas lipid-degradation functions were enriched in Cluster 4. (C) Amino acid and oligopeptide acquisition functions were most prominent in Clusters 1 and 2, whereas synthesis and degradation functions became more prominent in Cluster 3.

A similar pattern was observed among lipid transport and metabolism genes (Fig. 3B). Cluster 1 showed a mixed pattern, with 3 of 6 genes classified as lipid synthesis, 2 of 6 as lipid degradation, and 1 of 6 as lipid acquisition. Cluster 2 was dominated entirely by lipid synthesis (19 of 19 genes, 100%), and Cluster 3 remained synthesis-biased (7 of 8 genes, 88%), with a single degradation gene. In contrast, Cluster 4 shifted completely to lipid degradation (2 of 2 genes, 100%). Given the extensive membrane expansion accompanying RB growth and replication, the enrichment of lipid synthesis genes in Clusters 2 and 3 is consistent with the biosynthetic demands of the RB phase. The predominance of lipid degradation genes in Cluster 4 suggests membrane turnover and reorganization during EB maturation.

Genes involved in amino acid and oligopeptide transport and metabolism displayed a distinct developmental pattern (Fig. 3C). Acquisition functions accounted for 2 of 3 genes in Cluster 1 and 17 of 26 genes in Cluster 2, indicating substantial reliance on host-derived amino acids and peptides during early infection and RB establishment (8). In Cluster 3, acquisition declined to 8 of 24 genes, whereas synthesis and degradation accounted for 7 of 24 genes and 9 of 24 genes, respectively. No amino acid/oligopeptide genes fell into Cluster 4. These findings suggest that amino acid metabolism shifts from host-derived acquisition during early development to increased processing during active RB proliferation.

Together, these analyses indicate that the developmental cycle is accompanied by pathway-specific metabolic transitions rather than a single global metabolic switch. Energy metabolism shifts during RB establishment, lipid metabolism transitions from synthesis during RB growth to degradation during EB maturation, and amino acid/oligopeptide metabolism shifts from acquisition to increased processing during proliferation.

### Developmental distribution of transcriptional regulators

To gain insight into the regulatory architecture underlying developmental progression, we examined genes annotated in the transcription COG together with curated developmental regulators across the four transcriptional programs. Among curated developmental regulators, only Euo fell into Cluster 1 (Fig. 4) (Table S1). Euo is a well-characterized developmental regulator that represses late genes (14, 16, 17), consistent with a role in establishing an RB-competent developmental program while preventing premature activation of late developmental processes.

**Fig. 4.**
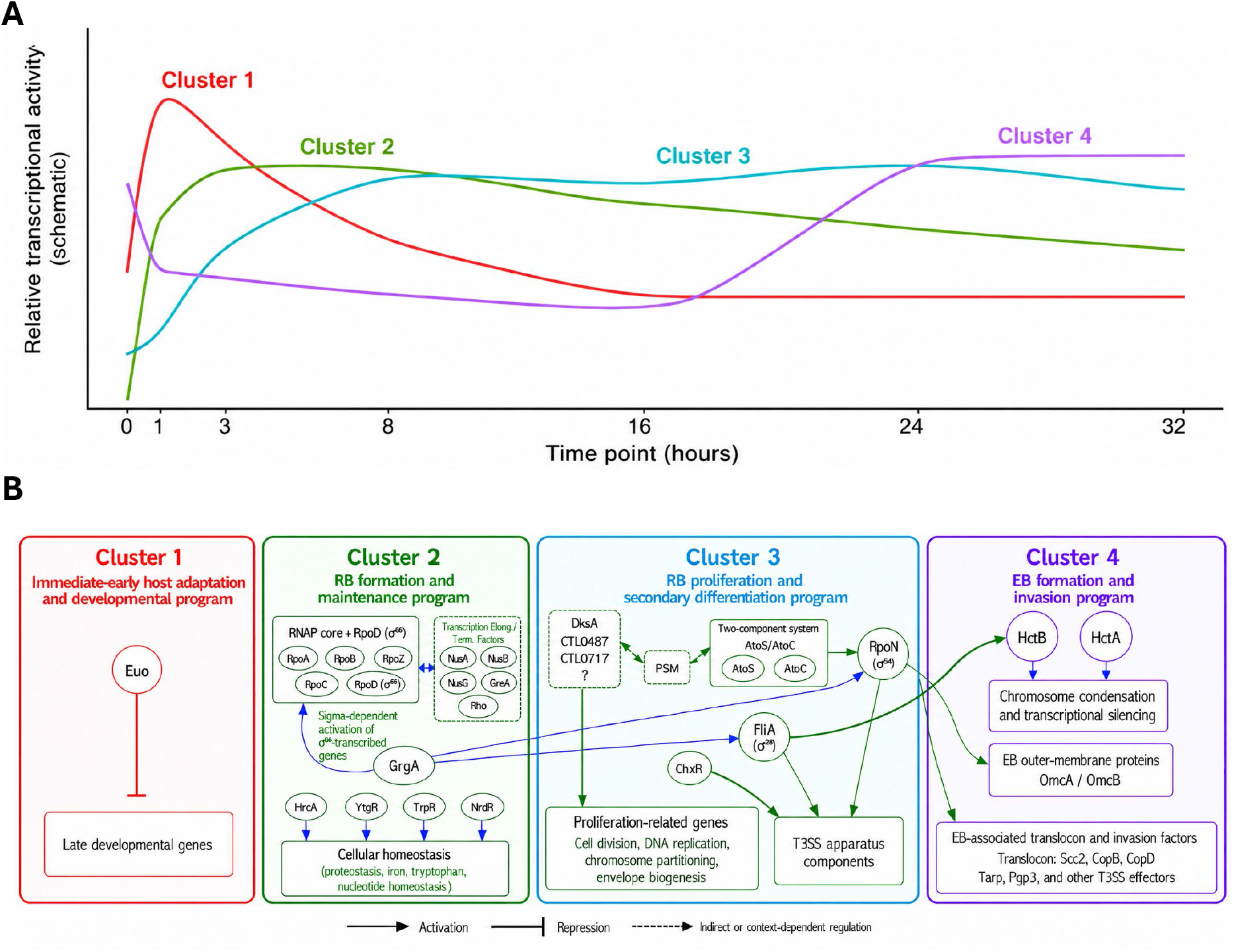
Regulatory and mechanistic features associated with the four coordinated transcriptional programs. (A) Schematic trajectories showing the relative timing, persistence, and overlap of the four transcriptional programs, informed by Fig. 2B. (B) Representative regulators and pathway components associated with each transcriptional program. Placement indicates cluster membership (Tables S1–S4), whereas connections depict regulatory relationships: arrows indicate activation, T-bars indicate repression, and dashed arrows indicate indirect or context-dependent regulation. RNAP, RNA polymerase; PSM, partner-switching mechanisms; T3SS, type III secretion system; Elong./Term., elongation and termination.

Cluster 2 contained the majority of genes involved in transcription initiation, elongation, and termination (Fig. 4). These included all major RNA polymerase structural subunits (RpoA, RpoB, RpoC, and RpoZ) (19), the primary sigma factor σ66 (RpoD), and multiple transcription elongation and termination factors, including NusA, NusB, NusG, GreA, and Rho (Table S2). This cluster also contained the repressors HrcA, YtgR, TrpR, and NrdR, which govern cellular functions essential for physiological fitness, including proteostasis, iron homeostasis, tryptophan metabolism, and deoxyribonucleotide synthesis (20–26). GrgA also fell into Cluster 2 (Table S2). Unlike the repressors in this cluster, GrgA functions as a global transcriptional activator required for EB formation (27); its placement within the RB formation and maintenance program is consistent with its contribution to optimal RB growth (27).

Cluster 3 contained several regulators implicated in developmental responsiveness and environmental adaptation, including the alternative sigma factors σ54 (RpoN) and σ28 (FliA), the *Chlamydia*-specific regulator ChxR, the DksA-family protein, and the AtoS/AtoC two-component regulatory system (28, 29) (Fig. 4, Table S3). Notably, AtoS/AtoC regulates transcription through the σ54 holoenzyme, linking two major regulatory components within this cluster (30). σ54- and σ28-dependent transcription has been linked to late gene expression required for EB formation (30–32). ChxR has also been implicated in late developmental regulation, as proteomic analyses of a *chxR* null mutant revealed reduced abundance of multiple type III secretion system effectors (28). Together, these observations suggest that Cluster 3 contains specialized regulatory circuits associated with developmental progression and deployment of virulence-associated functions during the proliferative phase.

Cluster 4 contained the histone-like proteins HctA and HctB, which are strongly associated with chromosome condensation and EB maturation (33) (Fig. 4). This cluster also contained CTL0463, a member of the YqgE/AlgH protein family (Table S4). Although the function of CTL0463 in *Chlamydia* remains unknown, its coexpression with HctA and HctB suggests that it may participate in late developmental processes.

## Discussion

In this study, we used high-coverage RNA-seq and unsupervised clustering of temporal expression profiles to refine the developmental transcriptomic landscape of *C. trachomatis*. Four coordinated transcriptional programs emerged across the cycle, revealing shifts from immediate host adaptation to RB formation and maintenance, RB proliferation and secondary differentiation, and EB formation and invasion. These programs should be viewed as coordinated and partially overlapping developmental priorities rather than mutually exclusive modules.

The principal conceptual advance is the distinction between transient immediate-early expression and sustained early expression. Our previous study established extensive transcriptional activation during the first hour of infection (8); the present analysis resolves how early-activated expression develops over the subsequent cycle. The separation of Clusters 1 and 2 distinguishes a transient response associated with the establishment of the intracellular niche from a more sustained program supporting the development and maintenance of biosynthetically competent RBs. Thus, the timing of initial activation and the subsequent expression trajectory provide complementary information about a gene’s developmental contributions.

The different trajectories of Clusters 1 and 2 may also reflect shifts in the allocation of limited translational capacity. Our previous measurements suggested that 23S rRNA availability could constrain ribosome assembly during the immediate-early phase, even as rRNA synthesis and expression of translation-associated genes increased (8, 34). Against this background, the transient Cluster 1 trajectory may favor production of proteins needed for host adaptation, while the later peak and sustained expression of Cluster 2 genes may support expansion and maintenance of bacterial biosynthetic capacity. This temporal organization suggests that changes in transcript availability may influence competition for translational resources while the protein-synthesis machinery itself expands.

The Cluster 2 trajectory also provides a possible transcriptional context for reported morphological changes during RB development. Its expression peaks at approximately 3 hpi, before RB formation is complete, and declines thereafter, whereas RB size was reported to decrease progressively from 12 hpi onward (4). The earlier transcriptional peak may support the biosynthetic expansion needed for RB formation, with the subsequent decline accompanying a shift toward continued proliferation. The temporal relationship is compatible with a connection between changing biosynthetic needs and RB morphology, although it does not establish a direct correspondence between relative transcript abundance and cell size.

The distinction between Clusters 2 and 3 further indicates that the establishment of a biosynthetically competent RB state and the expansion of the RB population are connected but supported by different gene expression programs. Biosynthetic maintenance continues during proliferation, while the coordinated expression of replication-associated genes and developmental regulators in Cluster 3 suggests that population expansion is accompanied by preparation for secondary differentiation. The high expression of Cluster 3 genes from approximately 8 hpi onward is consistent with RB proliferation and preparation for secondary differentiation following completion of primary differentiation.

GrgA provides an instructive example of how TPM-based clustering and expression measurements normalized to bacterial genome copy number provide complementary information. In the TPM analysis, *grgA* clusters with the RB formation and maintenance program, indicating that its relative expression trajectory is coordinated with genes supporting biosynthetic capacity during RB establishment. This placement is consistent with the demonstrated role of GrgA in supporting optimal RB growth (27). Measurements of transcripts per genome and protein abundance, however, showed that GrgA expression peaks at approximately 18 hpi, shortly before substantial RB-to-EB conversion (27). This later peak is consistent with its demonstrated requirement for secondary differentiation and late-stage transcriptomic activation (27). TPM profiles identify the coexpression program with which a gene is associated, whereas transcript-per-genome measurements quantify its transcript abundance per bacterial genome. Together with protein measurements, these approaches place GrgA within an RB-associated coexpression program while revealing its later accumulation and contribution to EB formation. More broadly, measurements normalized to genome copy number can complement TPM-based trajectories by clarifying how relative transcriptional priorities relate to transcript abundance per genome.

The distinction between Clusters 3 and 4 extends this framework to late development. The Cluster 3 program supports secondary differentiation, understood here as the functional and regulatory reprogramming that prepares proliferating RBs for the RB-to-EB transition, while the Cluster 4 program supports EB formation and invasion, encompassing the later structural and compositional maturation of infectious EBs, host-cell exit, and preparation for invasion in the next developmental cycle. The distinction, therefore, captures different contributions to late development while recognizing that regulatory preparation and structural maturation are temporally connected and partially overlapping.

This distinction between developmental preparation and EB maturation raises the possibility that proliferating RBs acquire differentiation competence before engaging the regulatory processes that promote the transition. Sustained expression of Cluster 3 genes may maintain this competence. GrgA is required for activation of the late transcriptional program needed for EB formation, and its increasing abundance toward an approximately 18-hpi peak (27) suggests a mechanism for coordinating differentiation readiness with progression toward EB formation.

Collectively, these findings refine the developmental transcriptomic landscape of *C. trachomatis* by distinguishing early host adaptation from RB biosynthetic establishment and maintenance, and by separating proliferation-associated developmental preparation from later EB maturation and invasion competence. The integration of these trajectories with GrgA expression and functional evidence further suggests that differentiation competence may be established before the regulatory processes promoting EB formation are fully engaged. This framework provides a basis for investigating how coordinated transcriptional programs and the timing of regulator accumulation govern progression through the chlamydial developmental cycle.

## Materials and Methods

### *Chlamydia* and culture

*C. trachomatis* L2 (strain 434/BU) was originally purchased from ATCC (35) and has been maintained in our lab. The bacterium was grown using L929 cells and Dulbecco’s modified Eagle medium containing 4.5 g/liter glucose and 110 mg/liter sodium pyruvate and supplemented with fetal bovine serum (final concentration, 5%), gentamicin (20 μg/ml), and cycloheximide (1 μg/ml). EBs for this study were sequentially purified through ultracentrifugation using 35% MD-76 and 44%/40%/52% MD-76 gradients.

### RNA preparation

Total chlamydial and host RNA was prepared as previously described with modifications (23). For 0 and 1 hpi, two previously reported RNA-seq datasets generated at MOIs of 50 and 200 were combined (8), with three biological replicates per MOI at each time point. Samples collected at 3, 8, 16, 24, and 32 hpi were obtained at MOIs of 50, 5, 1, 1, and 1 inclusion-forming units per host cell, respectively, with three biological replicates per time point. Infections were established by replacing the overnight culture medium in 6-well plates containing L929 monolayers with fresh medium containing EBs at the specified MOIs. The plates were centrifuged at 900 × g at room temperature for 10 min, then washed 3 times with 100 µg/ml heparin in HBSS to remove free and cell-surface-bound EBs. Completion of the washes was defined as 0 hpi. Cells designated for the 0-hpi sample were lysed immediately with Tri Reagent (Millipore Sigma); cells designated for subsequent time points were incubated at 37 °C in a CO2 incubator and lysed at the specified times. RNA in Tri Reagent was purified by following manufacturer’s instructions. Contaminating DNA was removed through 2 rounds of DNase I-XT. Complete DNA removal was confirmed by lack of amplification of *ctl0631*. RNA concentration was determined using Qubit RNA assay kit (ThermoFisher). Aliquots of the DNA-free RNA samples were stored at -80 °C.

### RNA-Sequencing

RNA-seq was performed as described with minor modifications (23, 36). Briefly, total RNA integrity was assessed using a Fragment Analyzer (Agilent) before library preparation. Mouse and chlamydial rRNAs were depleted using the Illumina MRZE706 Ribo-Zero Gold Epidemiology rRNA Removal Kit, and mouse mRNA was depleted using oligo(dT) beads. RNA-seq libraries were prepared using the Illumina TruSeq stranded mRNA-seq sample preparation protocol, quantified, pooled for cBot amplification, and sequenced on an Illumina HiSeq 3000 platform using a 50-bp single-read module. Short reads were aligned to the *C. trachomatis* L2 434/Bu genome, including the chromosome (GCF_000068585.1_ASM6858v1) and pL2 plasmid (AM886278), and to the mouse genome (GCF_000001635.27) using TopHat2. Gene-level read counts were obtained using HTSeq. For each of the 0- and 1-hpi time points, chlamydial read counts from the MOI-50 and MOI-200 datasets were pooled before TPM calculation. TPM values were calculated using only chlamydial transcript counts; mouse transcript counts were excluded from the normalization denominator. The resulting TPM profiles were used for downstream temporal expression analyses (37–39).

### Gene ontology analysis

Gene ontology analysis was performed based on COG functional classification of the *C. trachomatis* proteome (Galperin et al., 2015) with modifications recently reported (27, 40). Genes in selected metabolic categories were further classified by predicted physiological role, and their distributions across the four temporal clusters were determined.

## Acknowledgements

This work was supported by grants from the National Institutes of Health (Grant #AI140167 and AI154305 to HF and AI182210 to GZ). We acknowledge William Liu (Rice University) for his participation in the analysis of metabolic gene classifications and cluster distributions presented in Fig. 3. RNA-Seq data were generated in the Genome Sequencing Facility, which is supported by UT Health San Antonio, NIH-NCI P30 CA054174 (Cancer Center at UT Health San Antonio) and NIH Shared Instrument grant S10OD030311 (S10 grant to NovaSeq 6000 System), and CPRIT Core Facility Award (RP220662).

